# From Modulated Noise to Natural Speech: the Effect of Stimulus Parameters on the Frequency Following Response

**DOI:** 10.1101/864934

**Authors:** Jana Van Canneyt, Jan Wouters, Tom Francart

## Abstract

Frequency following responses (FFRs) can be evoked by a wide range of auditory stimuli, but for many stimulus parameters the effect on FFR strength is not fully understood. This complicates the comparison of earlier studies and the design of new studies. Furthermore, the most optimal stimulus parameters are unknown. To help resolve this issue, we investigated the effects of four important stimulus parameters and their interactions on the FFR. FFRs were measured in 16 normal hearing subjects evoked by stimuli with four levels of stimulus complexity (amplitude modulated noise, artificial vowels, natural vowels and nonsense words), three frequencies (around 105 Hz, 185 Hz and 245 Hz), three frequency contours (upward sweeping, downward sweeping and flat) and three vowels (Flemish /a:/, /u:/, and /i:/). We found that FFRs evoked by artificial vowels were on average 4 to 6 dB SNR larger than responses evoked by the other stimulus complexities, probably because of (unnaturally) strong higher harmonics. Moreover, response amplitude decreased with stimulus frequency but response SNR did not. Thirdly, frequency variation within the stimulus did not impact FFR strength, but only when rate of change remained low (e.g. not the case for sweeping natural vowels). Finally, the vowel /i:/ appeared to evoke larger response amplitudes compared to /a:/ and /u:/, but analysis power was too small to confirm this statistically. Differences in response strength between evoking vowels have been suggested to stem from destructive interference between response components. We show how a model of the auditory periphery can simulate these interference patterns and predict response strength. Altogether, the results of this study can guide stimulus choice for future FFR research and practical applications.

## 1. Introduction

Frequency following responses (FFRs) are phase-locked neural potentials that reflect the periodicity of the evoking auditory stimulus. They represent neural processing from the cochlea to the inferior colliculus with minor contributions from the auditory cortex (Bidelman, 2018; Coffey et al., 2016, 2017). As brought up by Bidelman and Powers (2018), surprisingly little is understood about the basic characteristics of the FFR. In this study, we investigated the relation between parameters of the evoking stimulus and FFR strength.

Per definition, FFRs are evoked by an auditory stimulus. This stimulus can be chosen quite freely, as long as it has some periodicity for the auditory neurons to phase-lock to. This flexibility makes the FFR a versatile response with large application potential. For example, FFRs are applied in research on fundamental auditory processing, like perceptual organization (Yamagishi et al., 2016), temporal integration (Xu and Ye, 2015) and selective attention (Etard et al., 2019; Forte et al., 2017). In addition, FFRs have been used to study more holistic aspects of auditory perception, like vowel identification (Won et al., 2016), consonant recognition (Plyler and Ananthanarayan, 2001) and pitch processing (Krishnan et al., 2009, 2010).

Nevertheless, the large amount of stimulus options is also a source of problems. For instance, it is not easy for researchers to decide what stimulus to use. Many applications benefit from a stimulus that evokes large responses in the majority of the population. This is particularly true in clinical context where fast and robust response measurement is key. At the moment, it is not known what the most optimal stimulus characteristics are and therefore stimulus choice is often based on past studies and intuition. Moreover, when comparing studies using the FFR, it is often not clear whether differences in results are caused by experimental parameters, subject factors or stimulus parameters. Without understanding the effect of stimulus parameters on the FFR, any comparative conclusion is unreliable.

One important stimulus parameter that differs widely between studies is stimulus complexity. Many studies use speech(-like) stimuli, e.g. artificial vowels (Krishnan, 2002, 1999; Lehmann and Schönwiesner, 2014; Schoof and Rosen, 2016; Won et al., 2016; Xu and Ye, 2015), natural vowels (Galbraith et al., 1998; Aiken and Picton, 2006, 2008; Easwar et al., 2015), natural syllables (e.g. /da/ or /ba/) (Coffey et al., 2017; Jenkins et al., 2017; Russo et al., 2004), words (Galbraith et al., 1995, 1997), sentences (Choi et al., 2013), or, more recently, continuous speech (Etard et al., 2019; Forte et al., 2017; Reichenbach et al., 2016). However, much simpler periodic stimuli are commonly used as well. These simple stimuli are often unnatural, but they allow for precise manipulation of the acoustic cues. Examples are tonebursts (Clinard et al., 2010; Gardi et al., 1979; Glaser et al., 1976; Tichko and Skoe, 2017), pure tones (Gockel et al., 2015; Holmes et al., 2018), tone sweeps (Billings et al., 2019; Clinard and Cotter, 2015; Krishnan and Parkinson, 2000; Purcell et al., 2004) and (sinusoidally) amplitude modulated (AM) stimuli (Bidelman and Patro, 2016; Dimitrijevic et al., 2016; Van Canneyt et al., 2019). It is unclear whether findings for these non-speech stimuli can be generalized to responses evoked by speech stimuli. Moreover, it is debatable which level of stimulus complexity is more optimal. One might argue that less complex stimuli evoke larger responses as it is easier for the auditory system to process them. On the other hand, more natural speech(-like) stimuli might cause enhanced responses because the brain is optimized for processing speech.

Another important factor is stimulus frequency. FFRs are typically evaluated for one frequency or frequency contour. For pure tones or tone bursts, this is the tone frequency. For AM stimuli, this is often the modulation frequency. Finally, for speech stimuli, this is usually the fundamental frequency of the voice (f0) or one of its harmonics. In general, a decrease in response amplitude with increasing tone frequency is observed, although this relation has peaks and valleys (Moushegian et al., 1973; Batra et al., 1986; Tichko and Skoe, 2017). Similarly, response amplitude generally decreases with increasing modulation frequency (Rees et al., 1986; Purcell et al., 2004; Gransier et al., 2016), with a well-studied peak near 40 Hz. For speech stimuli, the effect of frequency has not been evaluated.

An interesting characteristic of the FFR, which separates it from the auditory steady-state response, is that the frequency of interest may change over time. Krishnan and Parkinson (2000) and Billings et al. (2019) investigated the effect of glide direction and rate of frequency change for FFRs evoked by sweeping pure tones. Additionally, Purcell et al. (2004) studied FFRs to sweeps of modulation frequency. In speech stimuli, it is natural for f0 to change over time. This phenomenon, called intonation, is an inherent part of tonal languages and in non-tonal languages it conveys expressive meaning.Krishnan et al. (2004) measured FFRs to different Mandarin lexical tones, but a thorough analysis of the effect of intonation on FFR strength in speech stimuli is lacking.

The syllables /da/, /ba/ and /ga/ are very commonly used to evoke auditory responses, as they allow to measure both the speech-evoked auditory brain stem response (speech ABR) to the plosive sound and the sustained FFR response to the vowel (King et al., 2002; Wible et al., 2004; Ribas-Prats et al., 2019). The preference for these specific consonant-vowel combinationsthat the vowel /i:/ evoked larger responses than all other English vowels. This indicates that even for the most classic FFR paradigms, stimulus optimization might still be possible.

To increase understanding of the relation between stimulus parameters and response strength, we investigated how the FFR is affected by the four stimulus parameters discussed above, i.e. stimulus complexity, frequency, frequency contour and vowel identity. The effects on three FFR measures were evaluated, i.e. response amplitude, noise amplitude and response signal-to-noise-ratio (SNR). Importantly, the effects of stimulus frequency and frequency contour were studied for both speech and non-speech stimuli with focus on frequencies in the range of the f0. We aim to create a frame of reference for comparing studies that use different stimuli and to help determine the most optimal stimulus parameters to evoke the FFR.

## 2. Methods

### 2.1. Subjects

FFRs were measured for 16 participants (13 females, 3 males) with ages ranging between 18 and 24 years old (mean = 20.9 years, standard deviation = 1.6 years). All participants were confirmed to have normal hearing (all thresholds < 25 dB HL) using pure-tone audiometry (octave frequencies between 125 and 8000 Hz). The experiments were approved by the medical ethics committee of the University Hospital of Leuven and all subjects signed an informed consent form before participating.

### 2.2. Stimuli

The stimulus protocol included 20 stimuli that vary in complexity, frequency, frequency contour and vowel identity (see Figure 1). The 4 levels of stimulus complexity, i.e. AM noise (AMN), artificial vowels (AV), natural vowels (NV) and words (W), range from simple auditory stimulation to natural speech. The AM noise was a 450 ms section of ICRA speech-weighted noise (Dreschler et al., 2001) to which sinusoidal amplitude modulation was applied in MATLAB R2016b (The MathWorks Inc., 2016). The natural vowels were recorded from a female Flemish speaker (fs = 44.1 kHz). The recordings were trimmed to a 450 ms section with steady amplitude, beginning and ending in a zero crossing. The artificial vowel (duration = 450 ms) was created with a Klatt synthesizer (Klatt, 1980) implemented in MATLAB based on 5 parameters: the frequency and the bandwidth of the first two formants and the f0. These parameters were set to the values extracted from the recording of the natural vowel /i:/, i.e. the f0 was 180 Hz and the first two formant frequencies were 300 Hz (bandwidth = 50 Hz) and 2320 Hz (bandwidth = 440 Hz). The word stimulus, i.e. /i:di:/ (duration = 780 ms), was taken from the Leuven Analytical Speech Test (LAST). Example wave forms, spectrograms and envelope spectra of each of the stimulus types are presented in Figure 2.

**Figure 1:**
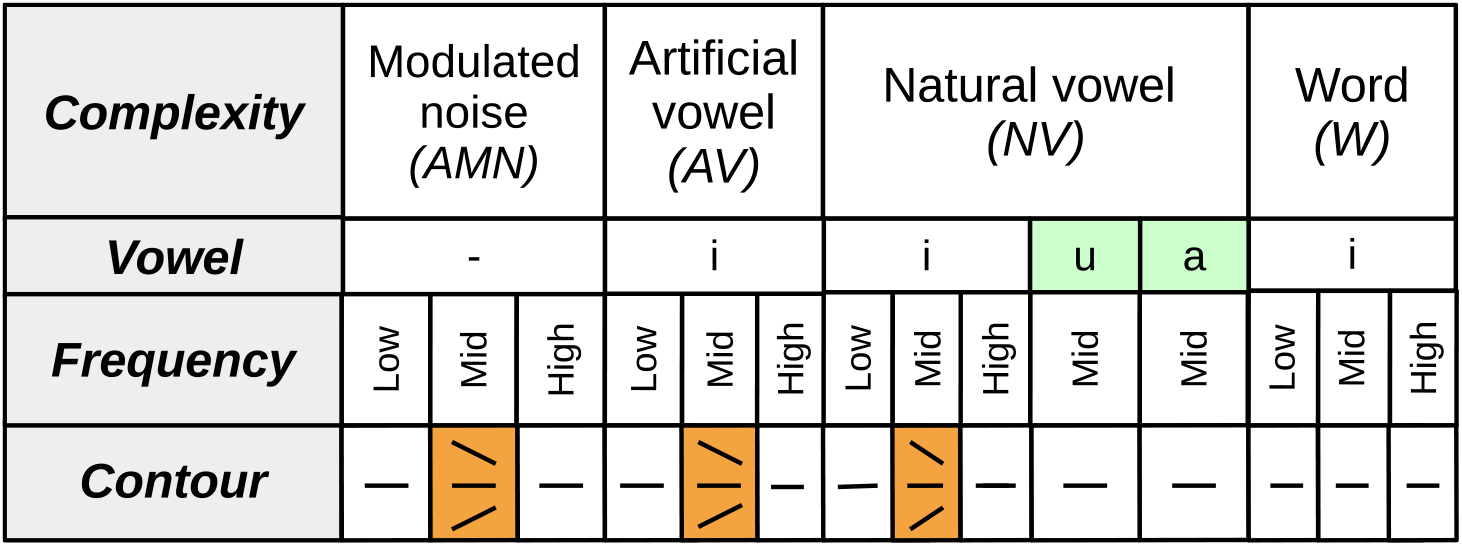
Stimulus overview. The stimuli included for analysis of the effect of stimulus frequency contour are indicated in orange. The stimuli used to analyze the effect of vowel identity are indicated in green.

**Figure 2:**
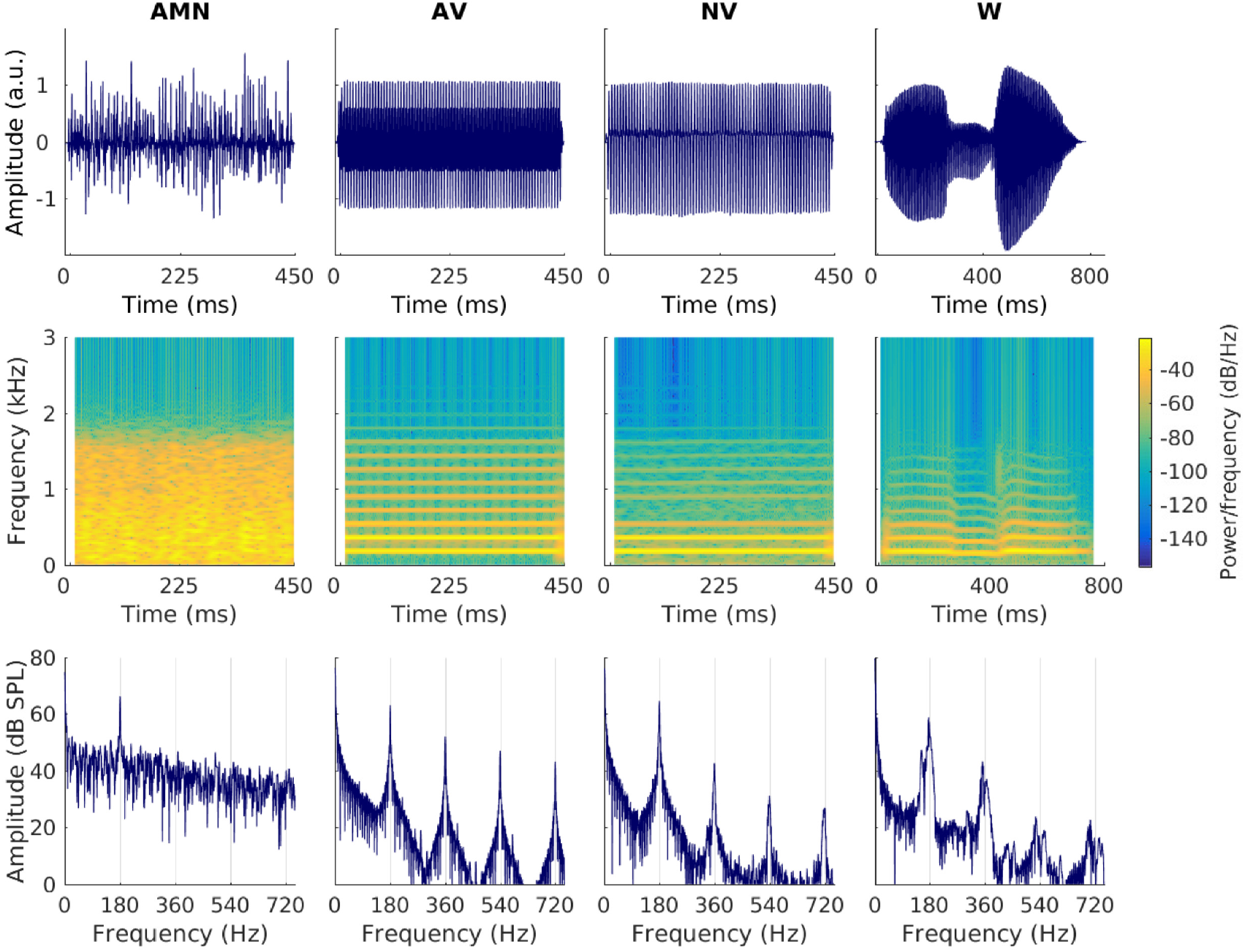
Example waveforms, spectrograms and envelope spectra of the stimulus types. AMN = amplitude modulated noise, AV = artificial vowel, NV = natural vowel, W = word. The f0 (or modulation frequency) for the stimuli in the figure is 180 Hz.

Each stimulus type was presented with three different frequencies: around 105 Hz (low), 180 Hz (mid) and 245 Hz (high). This is the modulation frequency for the AM noise, and the f0 for the other stimulus types. The mid and high frequency were chosen based on the recordings of the female talker, who was instructed to produce vowels with both her natural (f0 = 180 Hz) and a higher pitched voice (f0 = 245 Hz). The low frequency ensured that the full typical range of the adult f0 is represented, including the low f0 of male voices. The modulated noise and artificial vowels were created with the specified frequencies in MATLAB. The mid and high-frequency natural vowels were taken straight from the recordings. The low-frequency natural vowel was created by lowering the f0 of the mid-frequency stimulus to an average of 105 Hz using PRAAT (Boersma and Weenink, 2015). This operation did not distort the naturalness or intelligibility of the stimulus. Similarly, the pitch of the original word stimulus was adjusted towards the desired frequency values.

The AM noise stimuli, artificial vowels and natural vowels were presented with various frequency contours. Three contours were considered: upward, downward and flat. Sweeping natural vowels were produced by the speaker and then trimmed to 450 ms sections for which the frequency swept between 146 and 205 Hz (i.e. around the mid frequency defined above), either up or down (verified with PRAAT). The artificial vowels and AM noise stimuli were created with frequency contours changing linearly between the same frequencies.

All speech(-like) stimuli discussed up to now are based on the vowel /i:/, which Choi et al. (2013) found to provide the largest responses. However, FFR were also measured for the natural vowels /a:/ and /u:/, which together with /i:/ form the extremities of the Flemish triangle diagram. These vowels were also produced by the female speaker, with f0 equal to about 180 Hz. The frequencies of the first two formants were equal to 350 Hz and 915 Hz for /u:/ and to 800 Hz and 1600 Hz for /a:/.

All stimuli were resampled to a sampling frequency of 32 kHz and windowed with a 10 ms cosine attack and decay to reduce sharp on-and offsets in the stimulation. The stimuli were presented to the subject in random order, through an insert phone (ER3A, Etymotic) in the right ear using custom software (Hofmann and Wouters, 2010). Each stimulus was presented at 75 dBA for 600 repetitions with an interstimulus interval of 62 ms. Sound calibration was done using a 2 cc artificial ear (Bruel & Kjær, type 4152) and a sound level meter (Bruel & Kjær, type 2250).

### 2.3. Response measurement

FFRs evoked were measured with a 64-channel Biosemi ActiveTwo EEG recording system (fs = 8192 Hz). The 64 Ag/AgCl active scalp electrodes were placed on a cap according to the international standardized 10-10 system (American Clinical Neurophysiology Society, 2006). Two extra electrodes on the cap, CMS and DRL, functioned as the common electrode and the current return path, respectively. Subjects were seated in an electromagnetically-shielded sound-proof booth and watched a silent movie with subtitles. This encouraged the subjects to maintain the same passive listening state throughout the FFR measurements which lasted about 2 hours in total.

### 2.4. Response analysis

The EEG signals were processed with MATLAB. First, we averaged the signals of 12 electrodes located near the mastoids and the back of the head (CP5, CP6, P5, P6, PO3, PO4, O1, O2, PO7, PO8, TP7 and TP8) and referenced them to Cz, located at the top of the head. This electrode selection is the same as used in Gransier et al. (2016) and Van Canneyt et al. (2019). Then, the averaged signal was band-pass filtered between 2 Hz and 300 Hz (2nd order Butterworth filter). Next, 600 epochs were cut out of the filtered signal, with each epoch containing the response to one stimulus repetition. For stimulus frequencies above 80-100 Hz the neural processing delay is estimated at about 10 ms (Aiken and Picton, 2006; Forte et al., 2017). Therefore, for a 450 ms stimulus, we selected sections from 10 ms to 460 ms after the start of each stimulus. The epochs with the 5% largest peak-to-peak amplitudes were removed to reduce the amount of (muscle) artifacts.

The FFRs were analyzed using a Fourier Analyzer (Purcell et al., 2004; Aiken and Picton, 2006, 2008) with a reference estimated with the method of Aiken and Picton (2006, 2008). The Fourier Analyzer determines response strength along a particular frequency trajectory by complex multiplication of the response with orthogonal reference sinusoids matched to the instantaneous frequency of the stimulus. In case of artificial stimuli, this instantaneous frequency is known, but for natural stimuli it needs to be estimated. Aiken and Picton (2006, 2008) proposed a method to determine the instantaneous frequency based on the Hilbert transform of the stimulus and derivative of the instantaneous phase with respect to time. The instantaneous frequency is smoothed by filtering 3 times with a 1000-point moving average. This process introduces edge effects in the first and last 47 ms (1500 samples, fs = 32 kHz) of the instantaneous frequency contour, which were therefore cut both from the instantaneous frequency contour and from each response epoch. Word stimuli have a naturally slow attack and decay (see Figure 2) and in these areas of low amplitude the instantaneous frequency was imprecise as well. To make sure this did not influence the results, we discarded 60 ms at the start and 100 ms at the end of the frequency contour and response epochs for the word stimuli, leaving a stimulus of 620 ms. Finally, all frequency contours were down-sampled to the sampling frequency of the EEG, i.e. 8192 Hz, and used to create the orthogonal sinusoidal frequency references needed for the Fourier Analyzer. An overview of the estimated instantaneous frequency contours of all the stimuli in this study is presented in Figure 3.

**Figure 3:**
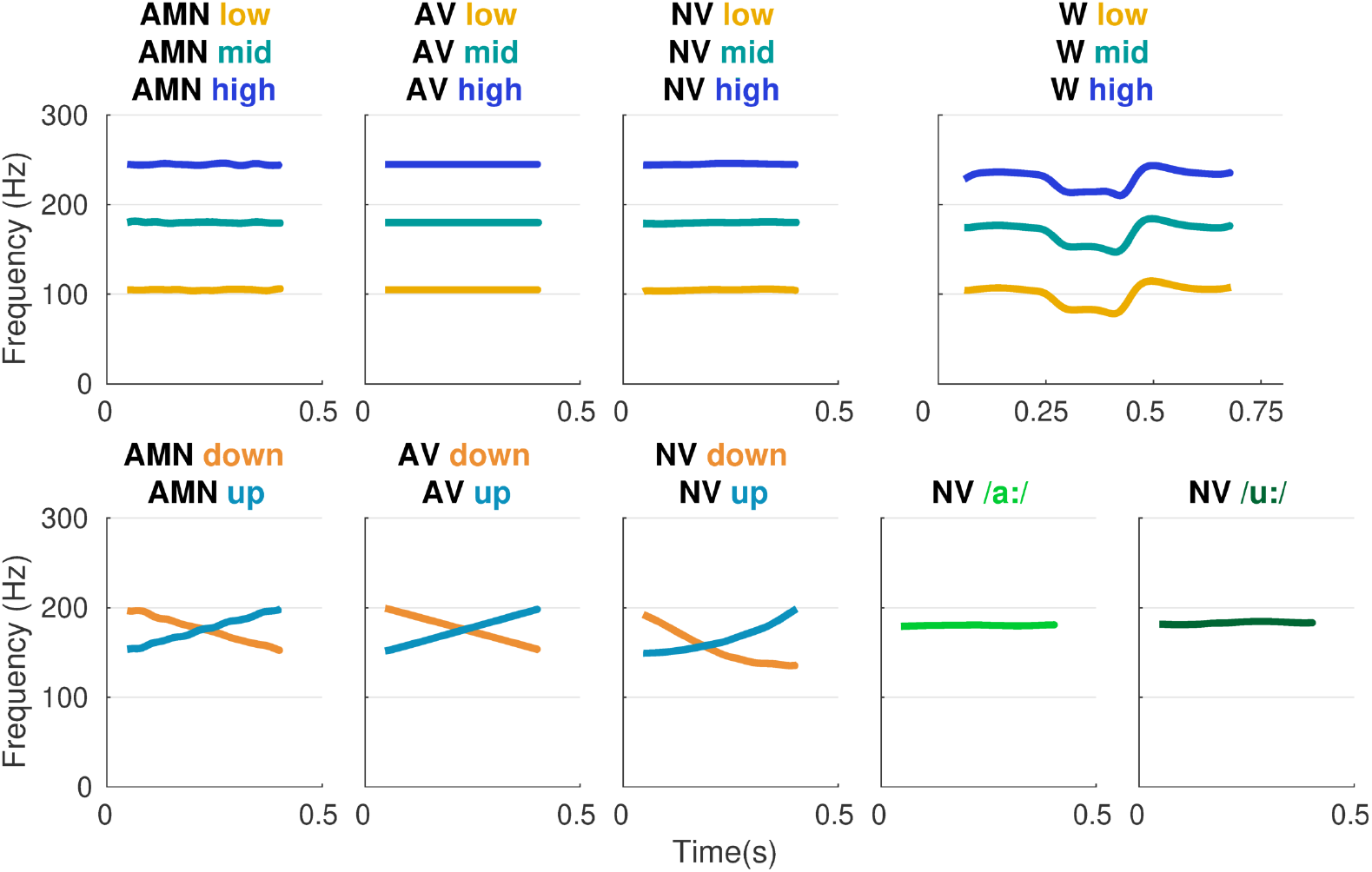
Overview of estimated instantaneous frequency contours. AMN = amplitude modulated noise, AV = artificial vowel, NV = natural vowel, W = word.

After applying the Fourier Analyzer and integrating along the epoch, the complex response amplitude per epoch is obtained. Then, the total response amplitude of the FFR is determined as the average magnitude of the complex response amplitude over epochs. The Hotelling T^2^ test (Hotelling, 1931) compares the total response amplitude with the noise amplitude, which is defined as the variance in response amplitude over epochs. This analysis provides both a significance value and a SNR estimate for each response measurement. A significance level of 5 % corresponds to a SNR of 4.8 dB, calculated with the method of (Dobie and Wilson, 1996). It is important to note that out of the 320 responses that were measured, 88 (27.5%) were not significant, i.e. the response could not be distinguished from the background noise. In section 4.2 it is explained that our selected stimulus frequency likely contributed to this. To avoid bias of excluding smaller responses, the non-significant responses were included in the analysis.

### 2.5. Statistical analysis

The effect of the stimulus parameters on the FFR was analyzed in R (R Core Team, 2018) with linear mixed models (LMM) (package lme4, version 1.1.17, Douglas et al. (2015)). We employed a random intercept per subject to account for the large inter-subject variability FFRs are known to have. Three LMMs were constructed for each of the stimulus parameters, i.e. one for each of the response outcomes: response amplitude (in nV), noise amplitude (in nV) and response SNR (in dB). For each stimulus parameter, we defined contrasts to test our hypotheses (see results section). Significance of the contrasts was evaluated with a t-test (significance level = 0.05) using Satterthwaite's method to estimate the degrees of freedom (Satterthwaite, 1946). Visual inspection of the residual plots from any of the models reported in this paper revealed no large deviations from homoscedasticity or normality.

## 3. Results

### 3.1. The effect of stimulus complexity

The effect of stimulus complexity was investigated for a subset of the data, including only the responses for stimuli with steady frequency contour and vowel identity /i:/ (see Figure 1). Response amplitude, noise amplitude and response SNR for the 192 responses (4 stimulus types x 3 frequencies x 16 participants) in this data set are visualized per stimulus type in Figure 4. The large whiskers of the boxplots indicate large intersubject variability (as expected for FFRs), confirming the need for a random intercept per subject in the linear mixed models. Three contrasts were defined. The first contrast considered a difference between responses evoked by non-speech and speech-like stimuli, i.e. between responses for modulated noise and artificial vowels. The second contrast was the difference between responses evoked by artificial speech and by natural speech, i.e. artificial vowels vs. both natural vowels and words. The third contrast compared responses for the natural vowels with responses for words.

**Figure 4:**
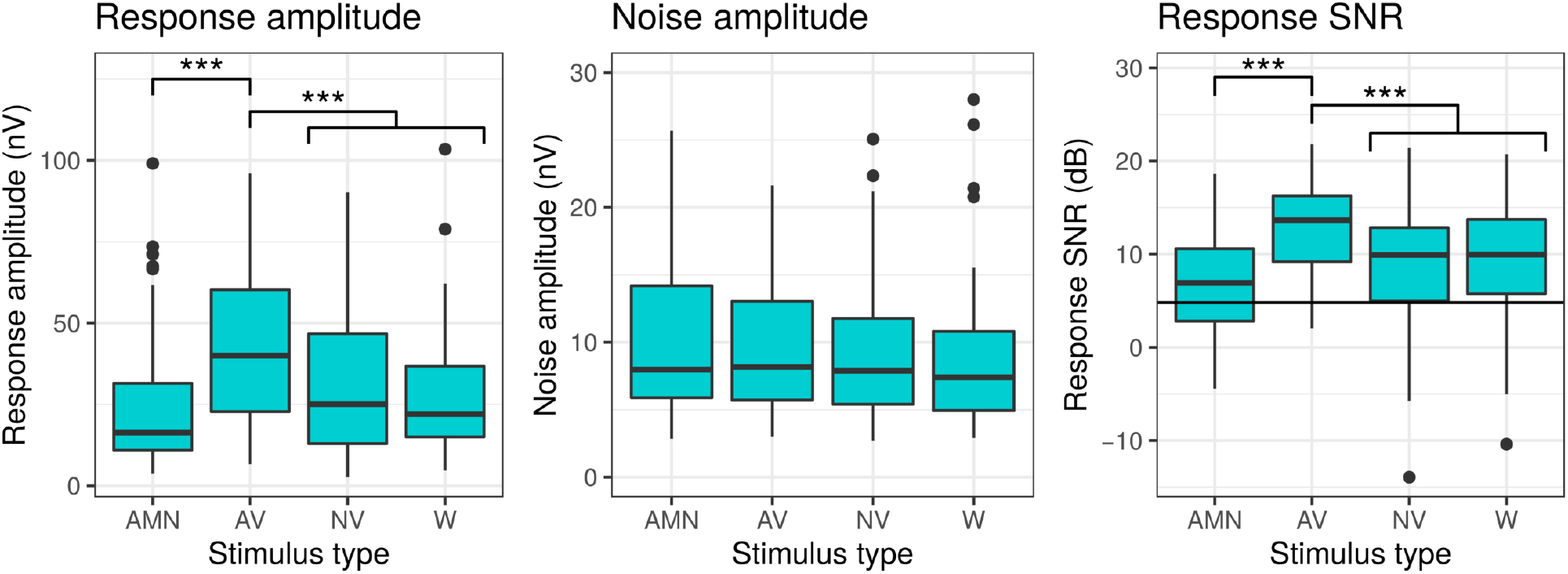
The effect of stimulus type on response amplitude, noise amplitude and response SNR. AMN = amplitude modulated noise, AV = artificial vowel, NV = natural vowel, W = word. Significance codes: *** < 0.001, ** < 0.01, * < 0.05. Only significant contrasts are indicated on the figure.

Response amplitude was significantly larger for artificial vowels than for modulated noise (p < 0.001), and compared to more natural speech stimuli, i.e. the natural vowels and the words (p < 0.001). The same pattern was found for response SNR (AMN vs. AV: p < 0.001; AV vs. NV+W: p < 0.001). There was no significant difference between natural vowels and words, both for response amplitude (p = 0.34), and for response SNR (p = 0.61). Noise amplitude was not significantly affected by stimulus type (AMN vs. AV: p = 0.98; AV vs. NV+W: p = 0.60; NV vs. W: p = 0.49). More details on this statistical analysis are presented in Table 1.

**Table 1:** Statistical analysis of the effect of stimulus complexity on the FFR

| Measure | Contrast | $\beta$ | df | t | p |
| --- | --- | --- | --- | --- | --- |
| Response amplitude | AMN vs. AV | -17.92 | 176 | -4.08 | <0.001 |
|  | AV vs. AMN + AV | 14.21 | 176 | 3.73 | <0.001 |
|  | NV vs. W | -4.23 | 176 | -0.96 | 0.337 |
| Noise amplitude | AMN vs. AV | 0.02 | 176 | 0.02 | 0.984 |
|  | AV vs. AMN + AV | 0.44 | 176 | 0.52 | 0.603 |
|  | NV vs. W | -0.67 | 176 | -0.69 | 0.493 |
| Response SNR | AMN vs. AV | -5.72 | 176 | -5.41 | <0.001 |
|  | AV vs. AMN + AV | 0.92 | 176 | 4.19 | <0.001 |
|  | NV vs. W | 0.53 | 176 | 0.51 | 0.614 |

Additionally, the effect of stimulus complexity was studied for low-, mid-and high-frequency stimuli separately. A detailed figure (Figure A.1) and overview of the statistical results (Table A.1, A.2 and A.3) of this analysis are available in the appendix. In general, the findings described above are true across stimulus frequencies. Remarkably, the advantage of artificial vowels over the other stimuli appears smaller for low-frequency compared to mid-and high-frequency stimuli.

### 3.2. The effect of stimulus frequency

The effect of stimulus frequency was studied for the same data set as described in the previous section. Response amplitude, noise amplitude and SNR are shown per stimulus frequency in Figure 5. Statistical analysis compared responses evoked by low and mid-frequency stimuli, as well as responses evoked by mid and high-frequency stimuli. The results indicate a significant decrease in response amplitude with increasing stimulus frequency (low vs. mid: p < 0.001; mid vs. high: p = 0.022). Results also show a decrease in noise amplitude with stimulus frequency (low - mid: p < 0.001; mid - high: p < 0.001). There was no monotonic relationship between response SNR and frequency. Mid-frequency stimuli evoked significantly smaller response SNRs than both low-frequency stimuli (p < 0.016) and high-frequency stimuli (p = 0.031). A more detailed overview of this statistical analysis is presented in Table 2.

**Figure 5:**
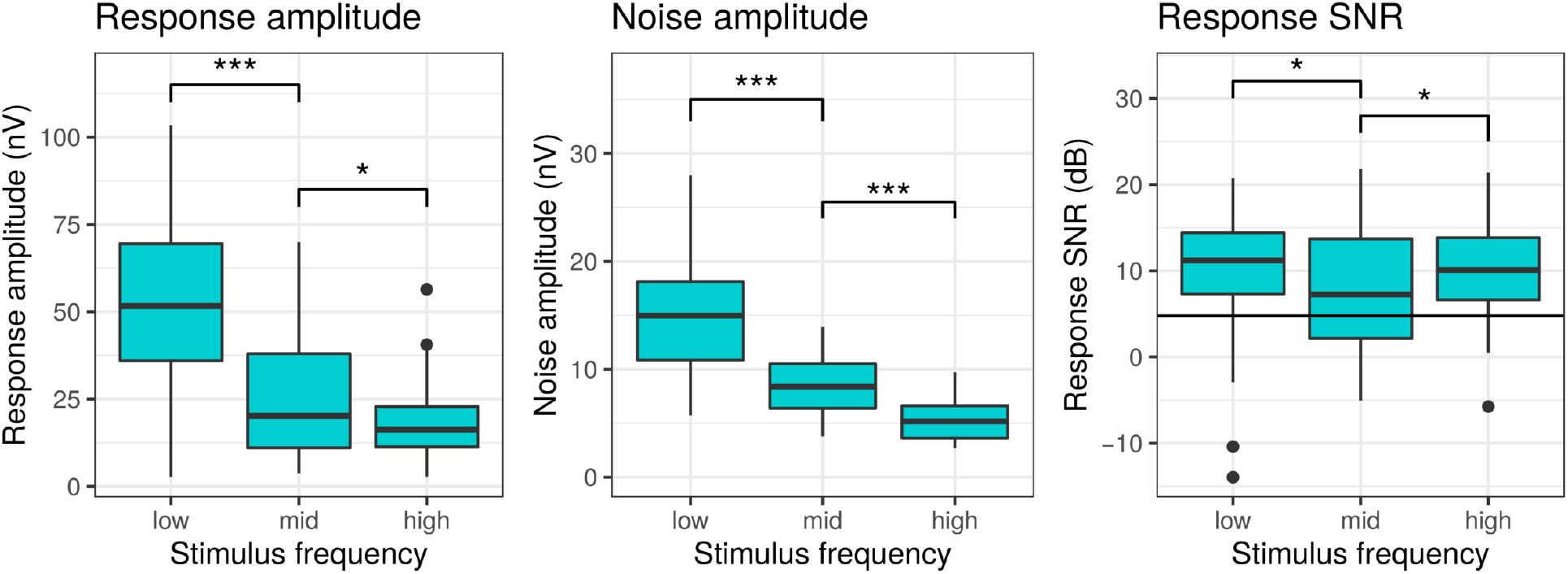
The effect of stimulus frequency on response amplitude, noise amplitude and response SNR. Low frequency ~ 105 Hz, Mid frequency ~ 180 Hz and high frequency ~ 245 Hz. Significance codes: *** < 0.001, ** < 0.01, * < 0.05.

**Table 2:** Statistical analysis of the effect of stimulus frequency on the FFR

| Measure | Contrast | $\beta$ | df | t | p |
| --- | --- | --- | --- | --- | --- |
| Response amplitude | low vs. mid | -27.48 | 176 | -9.33 | <0.001 |
|  | mid vs. high | 6.81 | 176 | 2.31 | 0.022 |
| Noise amplitude | low vs. mid | -6.46 | 176 | -15.64 | <0.001 |
|  | mid vs. high | 3.13 | 176 | 7.58 | <0.001 |
| Response SNR | low vs. mid | -2.38 | 176 | -2.44 | 0.016 |
|  | mid vs. high | -2.12 | 176 | -2.17 | 0.031 |

When analyzing the effect of frequency for each stimulus type separately, very similar patterns are observed. Remarkable findings are that a significant difference in response amplitude between the mid-and the high-frequency stimuli is only present for artificial vowels, not for the other stimulus types. Moreover, response SNR was significantly affected by stimulus frequency for AM noise, but not for any other stimulus type. In part, this could be due to the smaller statistical power of these analyses on smaller subsets of the data. Again, a detailed figure (Figure A.2) and overview of the statistical results (Table A.4, A.5 and A.6) of this analysis per stimulus complexity are available in the appendix.

### 3.3. The effect of stimulus frequency contour

The effect of stimulus frequency contour is studied for responses evoked by mid-frequency modulated noise, artificial vowels or natural vowels (indicated in orange on Figure 1). This gives rise to a data set of 144 responses (3 stimulus types x 3 contours x 16 subjects). The results are visualized in Figure 6. First, we compared responses evoked by flat and variable frequency contours (up or down). Results indicate no significant difference for response amplitude (p = 0.513) or response SNR (p = 0.287). Noise amplitude was significantly lower for responses evoked by flat intonation (p = 0.005). Secondly, responses evoked by a down-contour and an up-contour were compared. There was no significant difference for response amplitude (p = 0.941), noise amplitude (p = 0.942) or SNR (p = 0.917).

**Figure 6:**
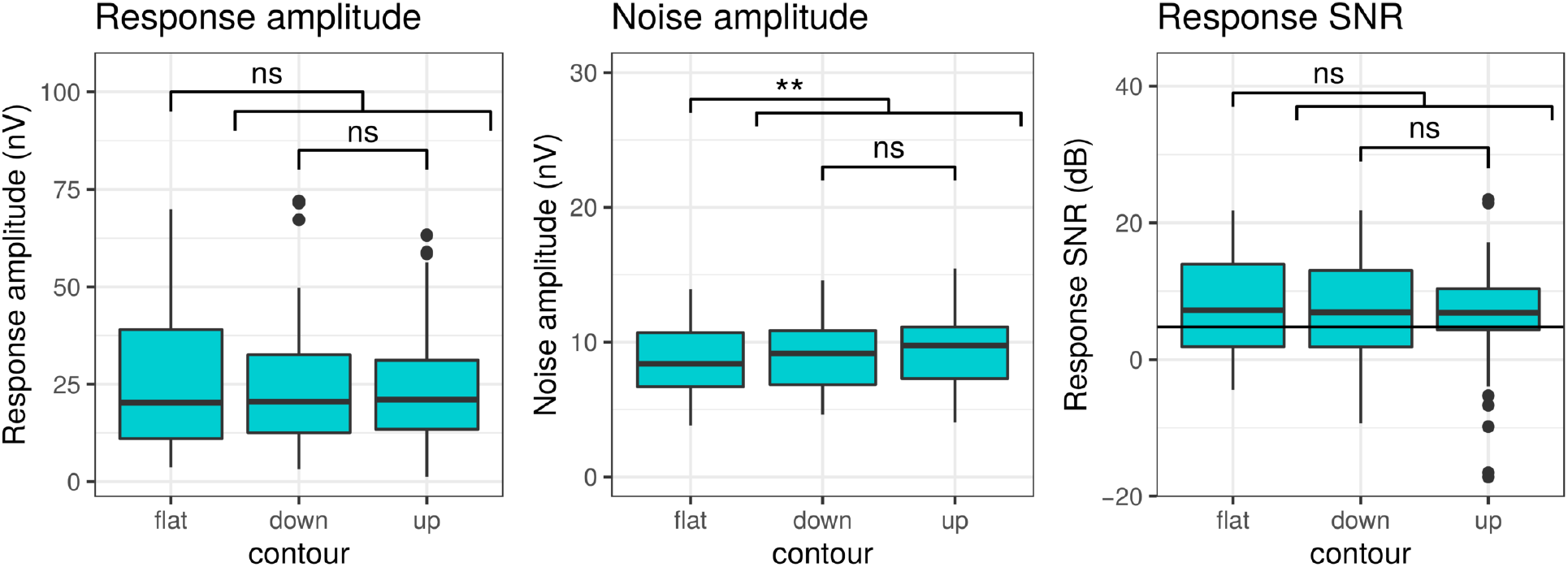
The effect of stimulus intonation on response amplitude, noise amplitude and response SNR. Significance codes: *** < 0.001, ** < 0.01, * < 0.05, ns = non-significant.

Once more, a more thorough analysis was performed to investigate the effect of stimulus frequency contour for each level of stimulus complexity separately. The corresponding figure (Figure A.3) and statistical data (Table A.7, A.8, A.9) are available in the appendix. Responses evoked by amplitude modulated noise and artificial vowels were not significantly influenced by frequency contour. In contrast, for natural vowels, the non-flat frequency contours evoked responses with significantly smaller response amplitude (p = 0.027) and SNR (p = 0.014), as well as larger noise amplitude (p < 0.001).

### 3.4. The effect of vowel identity

The effect of vowel identity is compared for mid-frequency natural vowels (see green on Figure 1). We compared the responses for /i:/ with responses evoked by /a:/ and with responses evoked by /u:/. Results (see Figure 7 and Table 4) show no difference in response amplitude between vowels (/i:/ vs. /u:/: p = 0.175; /i:/ vs. /a:/: p = 0.107). Noise amplitude was significantly larger for /i:/ than for /u:/ (p < 0.001) and for /a:/ (p = 0.038). Response SNR did not differ significantly between vowels (/i:/ vs. /u:/: p = 0.836; /i:/ vs. /a:/: p = 0.659).

**Figure 7:**
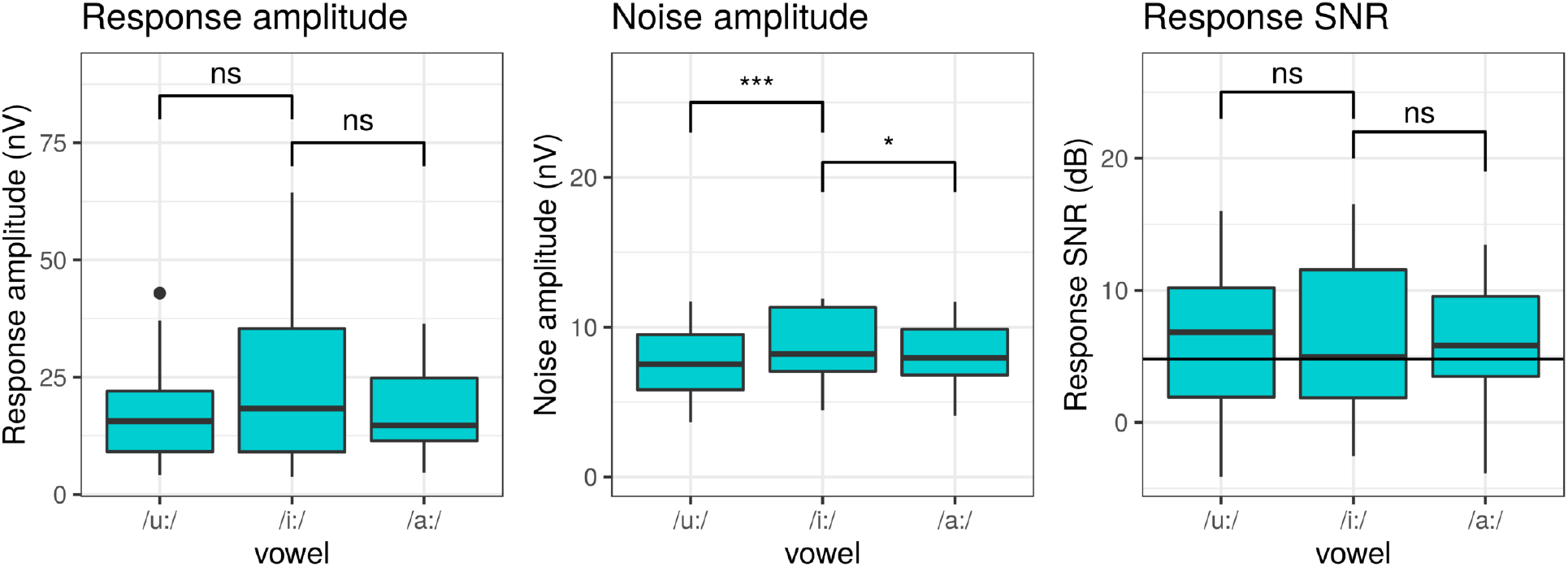
The effect of vowel identity on response amplitude, noise amplitude and response SNR. Significance codes: *** < 0.001, ** < 0.01, * < 0.05, ns = non significant.

**Figure 8:**
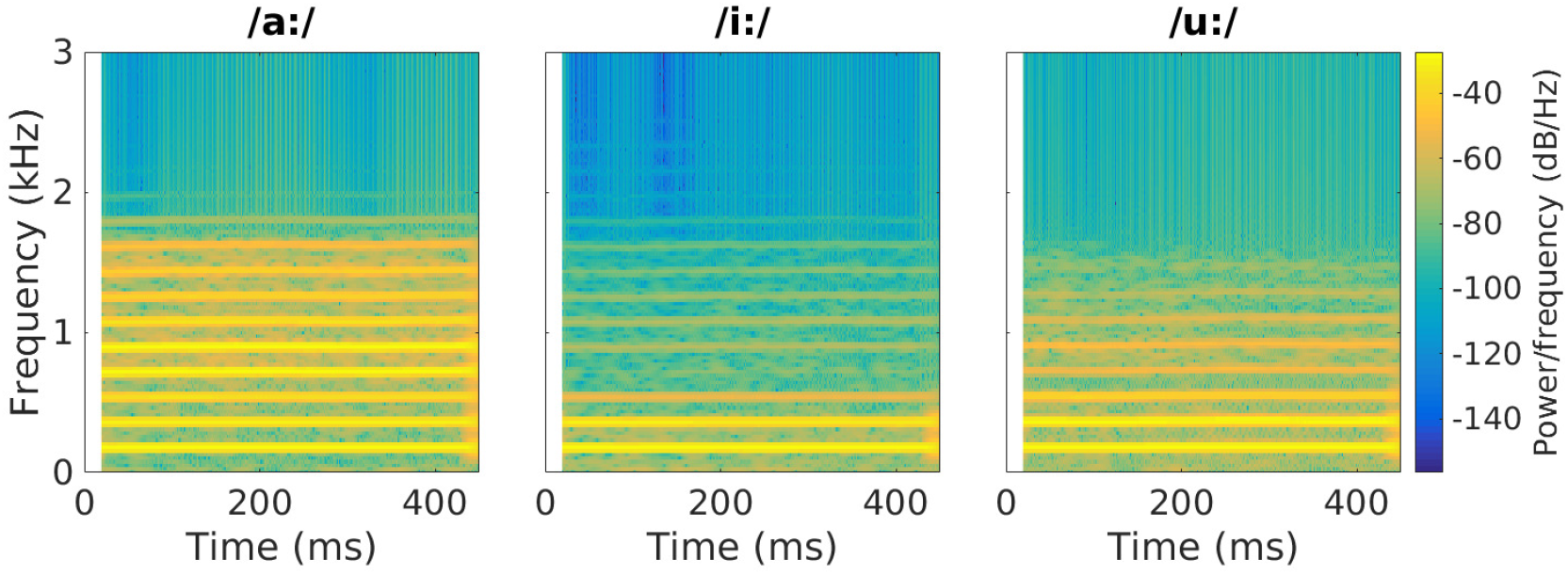
The effect of vowel identity on response amplitude, noise amplitude and response SNR. Significance codes: *** < 0.001, ** < 0.01, * < 0.05, ns = non significant.

**Table 3:**
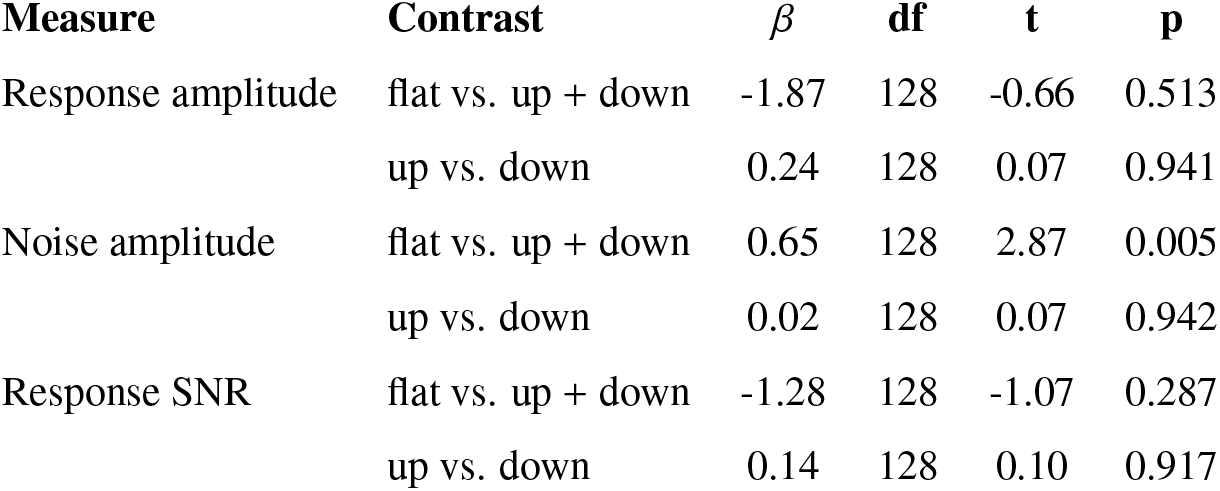
Statistical analysis of the effect of stimulus frequency contour on the FFR

**Table 4:** Statistical analysis of the effect of stimulus vowel identity on the FFR

| Measure | Contrast | $\beta$ | df | t | p |
| --- | --- | --- | --- | --- | --- |
| Response amplitude | /i:/ vs. /u:/ | 4.93 | 32 | 1.39 | 0.175 |
|  | /i:/ vs. /a:/ | 5.90 | 32 | 1.66 | 0.107 |
| Noise amplitude | /i:/ vs. /u:/ | 0.95 | 32 | 3.88 | <0.001 |
|  | /i:/ vs. /a:/ | 0.53 | 32 | 2.16 | 0.038 |
| Response SNR | /i:/ vs. /u:/ | 0.37 | 32 | 0.21 | 0.836 |
|  | /i:/ vs. /a:/ | 0.79 | 32 | 0.45 | 0.659 |

## 4. Discussion

To better understand the relation between characteristics of the evoking stimulus and FFR strength, we investigated the effect of 4 stimulus parameters on the response amplitude, noise amplitude and SNR of the FFR. The studied parameters were stimulus complexity, frequency, frequency contour and vowel identity. Below, the effects for each of these stimulus parameters will be discussed separately first. Then, the consequences with regards to stimulus choice for the FFR are reviewed.

### 4.1. The effect of stimulus complexity

Responses evoked by stimuli of increasing complexity and naturalness were compared. Results show that speech-like artificial stimuli evoked larger responses than both natural speech stimuli (either vowels or words) and less complex non-speech stimuli (AM noise). This effect was present both in terms of response amplitude and SNR. Overall, there was no significant difference in response amplitude or SNR between responses evoked by natural vowels and words. Moreover, noise amplitude was not significantly affected by stimulus type. This was expected because background EEG noise depends mostly on stimulus frequency and measurement duration.

The benefit of the artificial vowel over the other stimuli is likely driven by two factors. The first factor is spectral bandwidth. For both AM noise and the speech(-like) stimuli, the frequency of interest is present as an amplitude modulation across all frequency components of the stimulus. Therefore all primary neurons across this range will synchronize their firing patterns to this frequency of interest and contribute to the FFR (Aiken and Picton, 2006; Laroche et al., 2013). Moreover, it has been shown that neural activity in mid-to high-frequency regions of the cochlea is more important for FFR generation than activity in low-frequency regions (Dau, 2003; Vanheusden et al., 2019). Consequently, stimuli with high level components across a broad frequency range will cause more neurons to fire strongly and synchronously to the frequency of interest, evoking larger FFRs. Figure 2 shows how the AM noise and artificial vowel in this study have strong spectral components across a broader frequency range than the natural vowel and the word. The second factor is harmonic strength. Higher harmonics tend to fall with two or more in the same auditory filter (unresolved harmonics), and the summed response to multiple harmonics is modulated with the frequency difference between the harmonics, i.e. the f0 (Choi et al., 2013; Krishnan and Plack, 2011; Oxenham, 2008). This way strong unresolved higher harmonics can contribute to larger FFRs at f0 (Jeng et al., 2011; Laroche et al., 2013). Speech(-like) stimuli have higher harmonics whereas AM noise does not. Thus, the artificial vowel could be the most optimal stimulus because it combines high-level broad band energy with strong higher harmonics.

One limitation of this study is that the natural word stimulus did not have the same length as the other stimuli. The FFR measures evoked by words were based on 600 repetitions of a 780 ms stimulus, of which 620 ms were analyzed. In contrast, the FFR measures for natural vowels included 600 repetitions of a 450 ms stimulus of which 356 ms were analyzed. Therefore more data was averaged for the word condition, which reduces response noise. Figure 4 indeed shows a slightly reduced noise level for word stimuli compared to the other stimuli, however it was not significant. To verify whether this biased our results, we compared response strength for 345 repetitions of the word stimuli, with 600 repetitions of the natural vowel stimuli, which equals about 214 seconds of data for each condition. However, even with this modification, there was no significant difference between responses evoked by natural vowels and by words.

### 4.2. The effect of stimulus frequency

It is generally accepted that FFRs to modulated stimuli decrease in response amplitude for increasing modulation frequency (Rees et al., 1986; Picton et al., 2003; Purcell et al., 2004; Gransier et al., 2016), because precision of neural phase-locking is limited and the more rapid modulations are encoded progressively worse in the neural response. The results of this study confirm this decreasing trend and show that it is also applicable to speech(-like) stimuli. This indicates that speech(-like) stimuli with lower f0, i.e. typically male voices, will provide larger response amplitudes. However, evoked responses are best evaluated using response SNR so the recording noise is taken into account. The spectrum of background EEG noise is known to have a inversely proportional relationship with frequency (Cohen et al., 1991; Picton et al., 2003; Purcell et al., 2004), which is reflected in our results. Because both response amplitude and noise amplitude decrease with frequency, response SNR is relatively steady over frequencies. This has the advantage that the we can obtain qualitative responses with both male and female voices (unlike what was suggested based on amplitude alone).

It has been shown that the generally decreasing relation between response amplitude or SNR and frequency has many peaks and valleys (Gransier et al., 2016; Kuwada et al., 2002; Purcell et al., 2004; Purcell and John, 2010; Tichko and Skoe, 2017), in part because of destructive and constructive interference between various FFR sources (Tichko and Skoe, 2017). In this study, it was found that response SNR was significantly lower for the mid-frequency stimuli compared to the low-and high-frequency stimuli, indicating a valley near the mid-frequency, i.e. 180 Hz. The majority of the responses in this study were evoked with this frequency, explaining why so many were not-significant. This shows that frequency choice can have large impact on a study. In addition, the frequency values at which peaks and valleys occur are also found to be individually variable. This means the most optimal frequency to evoke the FFR is likely subject dependent.

### 4.3. The effect of stimulus frequency contour

Responses evoked by stimuli with a varying frequency contour did not differ significantly in response amplitude or SNR from responses evoked by stimuli with a flat frequency contour. This indicates that the FFR represents varying frequency equally well as steady frequency. Noise amplitude was somewhat larger for stimuli with varying intonation compared to stimuli with flat intonation. This is logically explained by the fact that the stimuli with flat intonation had a fixed frequency of 180 Hz, whereas the stimuli with variable frequency also contained lower frequencies for which noise levels are higher.

In contrast with the general conclusion, natural vowels with variable frequency contour did evoke significantly smaller responses than steady natural vowels. These contradictory results can be likely be explained by differences in the shape of the frequency contour (see Figure 3). The natural vowels were taken from recordings and therefore, in contrast with the other manually created stimuli, did not have a perfectly linear frequency contour (see Figure 3). The rate of change of the manually created stimuli (and the stimuli of Purcell et al. (2004)) was steady and relatively slow, i.e. 131 Hz/s (and 5.3-8.7 Hz, respectively). In contrast, the contours of the natural vowels have areas of rapid frequency change. The reduced amplitude and SNR for the natural vowels might be due to poor neural coding of these faster frequency changes. In studies that evoke FFRs with sweeping pure tones, less robust FFRs are found for higher rates of change, i.e. in the range of 900-6600 Hz/s (Billings et al., 2019; Clinard and Cotter, 2015). Another possibility is that the reference for the Fourier Analyzer represented these rapid frequency changes less precisely, impairing response analysis.

The directional (a)symmetry in FFR strength for pure tones with upward or downward sweeping frequency (with similar rate of change) is heavily discussed in literature. There are studies that find or imply larger neural synchrony for rising tones compared to falling tones (Collins and Cullen, 2005; Krishnan and Parkinson, 2000; Maiste and Picton, 1989), however there are also studies that do not find a difference (Billings et al., 2019; Arlinger et al., 1977; Clinard and Cotter, 2015; Elliott et al., 2005). The (a)symmetry in FFR strength for modulated or speech stimuli is much less studied. Purcell et al. (2004) found no significant difference between upward or downward sweeping modulation frequency in AM noise stimuli. The results of the present study confirm this and show that this finding can be generalized to speech-like stimuli.

### 4.4. The effect of vowel identity

We found no significant differences in response amplitude or SNR between responses evoked by /i:/ and /u:/ or by /i:/ and /a:/. Based on Figure 7, it seems that /i:/ might provide slightly larger response amplitudes than the other vowels, confirming the results of Choi et al. (2013), but this study lacks the statistical power to confirm this. Interestingly, the advantage of /i:/ over the other vowels, seems to contradict the theory explained in section 4.1. The spectrograms in Figure 2 show that /a:/ has broader high level activity and stronger harmonics than the other two vowels, suggesting that /a:/ would evoke the largest responses.

An explanation for this contradiction is found within the recent work of Easwar et al. (2018a,b). These studies show that there is destructive interference between responses initiated by distinct stimulus frequencies as they are measured at the scalp. The destructive interference occurs because of phase differences between the components, caused by earlier activation of high-frequency areas compared to low-frequency areas on the basilar membrane (John and Picton, 2000; Picton et al., 2003). The response to /a:/ is predominantly formed by a low-frequency component (from the first two harmonics) and a high frequency component from the fourth and fifth harmonic (enhanced by the first formant at 850 Hz). Following the formula proposed by Schoonhoven et al. (2002), there is 3 ms time delay between these components. With a fundamental frequency of 180 Hz, this translates to a phase difference of about 160° degrees, indicating considerable destructive interference takes place. In a similar vein, but to a lesser degree because the harmonics are less strong, responses to the vowel /u:/ will experience destructive interference between activity evoked by the first harmonics (enhanced by the first formant at 350 Hz) and harmonics near the second formant (915 Hz). The response to the vowel /i:/ does not suffer from this effect as the first formant is very low, i.e. 304 Hz, and therefore coincides with the strong first two harmonics, and the second formant, i.e. 2320 Hz, is sufficiently high to not cause strong harmonics.

To predict to what degree a stimulus will suffer from these destructive interaction effects originating in the cochlea, one can employ a model of the auditory periphery, e.g. the model of Bruce et al. (2018). Using the technique explained in Van Canneyt et al. (2019), we obtained simulated responses for each vowel stimulus. As shown in that study, the modulation depth of the simulated response for a particular stimulus, is a good predictor of the response SNR of the FFR evoked by that stimulus. The median modulation depth of the simulated responses was 790, 1700 and 846 spikes per time bin (3.2 ms), for the vowels /a:/, /i:/ and /u:/ respectively. These simulated results match the pattern observed in this study, confirming that these type of model simulations can be used to verify the effect of destructive interaction effects on the FFR.

Finally, responses for /i:/ had significantly larger noise amplitude compared to the other vowels. This is remarkable, as noise amplitude is expected to depend predominantly on stimulus frequency and measurement time, and both of these factors were equal across vowels. Since response amplitude and noise amplitude varied similarly over vowels, the ratio between the two measures stayed relatively constant and response SNR did not differ as much between vowels.

### 4.5. The most optimal stimulus to evoke the FFR

To obtain the largest possible response SNR, one should consider using an artificial vowel (or other stimulus with strong higher harmonics). The advantage that can be gained when using artificial vowels instead of modulated noise is about 5.7 dB SNR. Similarly, the advantage of using artificial vowels over natural speech is around 4 dB SNR. Artificial speech stimuli are an interesting compromise from a methodological point of view, i.e. they can be created as desired with full control over the stimulus and yet, they are relevant to speech understanding in real life. Relevance for day to day communication is especially important for applications like evaluation of hearing aid fitting (Choi et al., 2013; Scollie and Seewald, 2002).

Regarding the frequency of the stimulus, low-frequency stimuli are generally more optimal when response amplitude is considered, but not for response SNR. The relation between frequency and SNR is highly non-monotonous and idiosyncratic, hindering the selection of an optimal frequency. Furthermore, frequency can vary over time without compromising response strength, as long as rate of change is slow and steady. Moments of rapid frequency change, however brief, can weaken the FFR response.

An important condition to obtain large response amplitudes is that strong harmonics are spectrally located such that destructive interference is minimal. For some vowels, the formants enhance harmonics in a way that leads to stronger destructive interference compared to other vowels, with noticeable effects on response amplitude. In this context, the vowel /i:/ might be preferred over other vowels to evoke large FFRs. Importantly, the fundamental frequency of the stimulus influences the spectral positioning of harmonics and phase difference between responses components. Therefore, apart from possible formant locations, the fundamental frequency should also be taken into account when evaluating interference patterns. Simulations from a model of the auditory periphery (as done in Van Canneyt et al. (2019)) can be used to compare and predict the interference effects of different stimuli. As noise amplitude seems to be affected in a similar way, this matter might be of less importance for obtaining large response SNR.

Naturally, obtaining the largest response SNR is often not the main goal and some aspects of the evoking stimulus will be determined by the research question. But researchers are often left with at least some stimulus choices, and optimizing those can make a considerable difference in the amount of significant responses that are obtained. Besides stimulus optimization, the emerging knowledge of the effect of stimulus parameters on the FFR aids with study design, result interpretation and study comparison.

## A. Appendices

### A.1. The effect of stimulus complexity per stimulus frequency

**Figure A.1:**
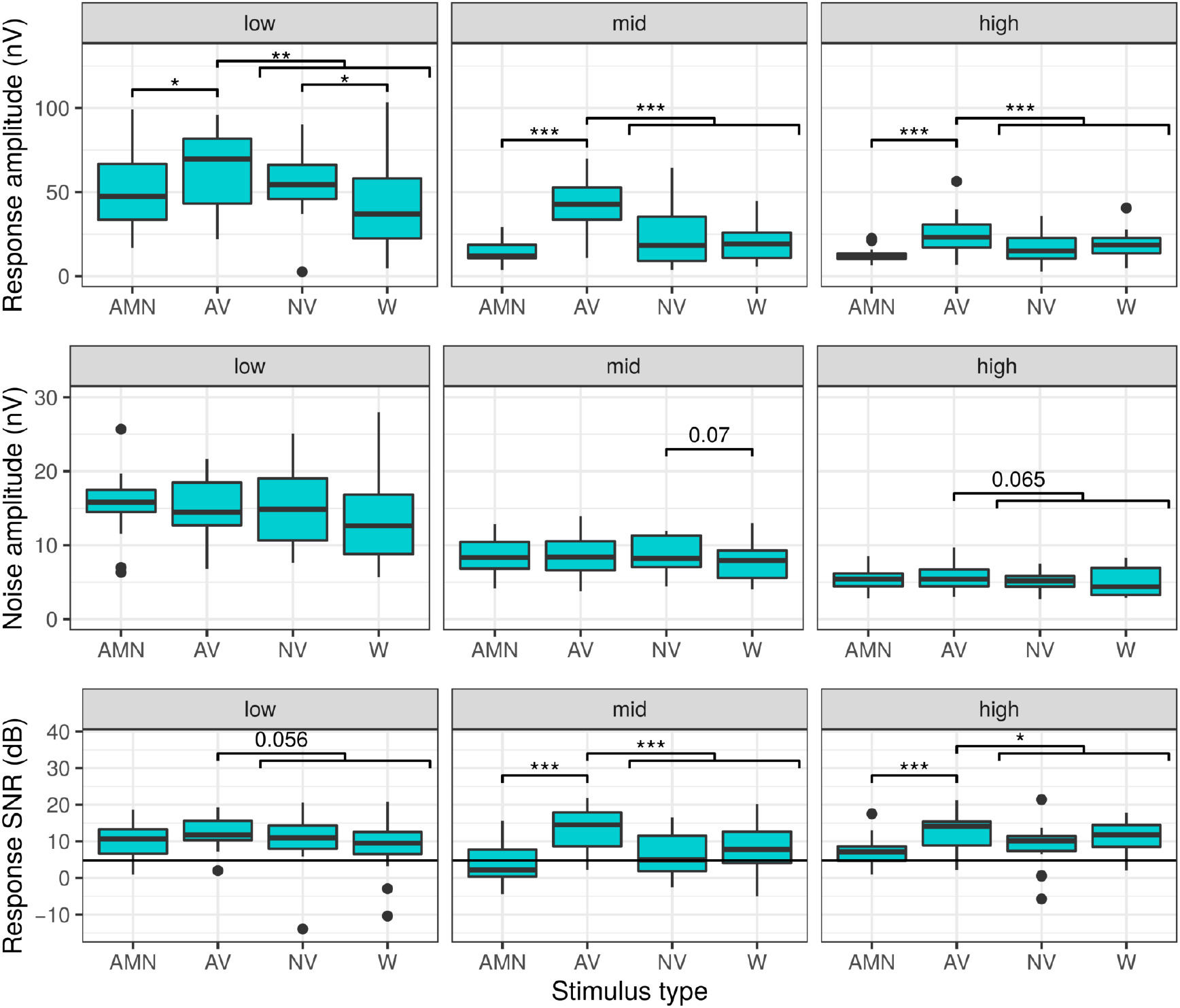
The effect of stimulus complexity on response amplitude, noise amplitude and SNR per stimulus frequency. AMN = amplitude modulated noise, AV = artificial vowel, NV = natural vowel, W = word. Significance codes: *** < 0.001, ** < 0.01, * < 0.05, ns = non significant.

**Table A.1:**
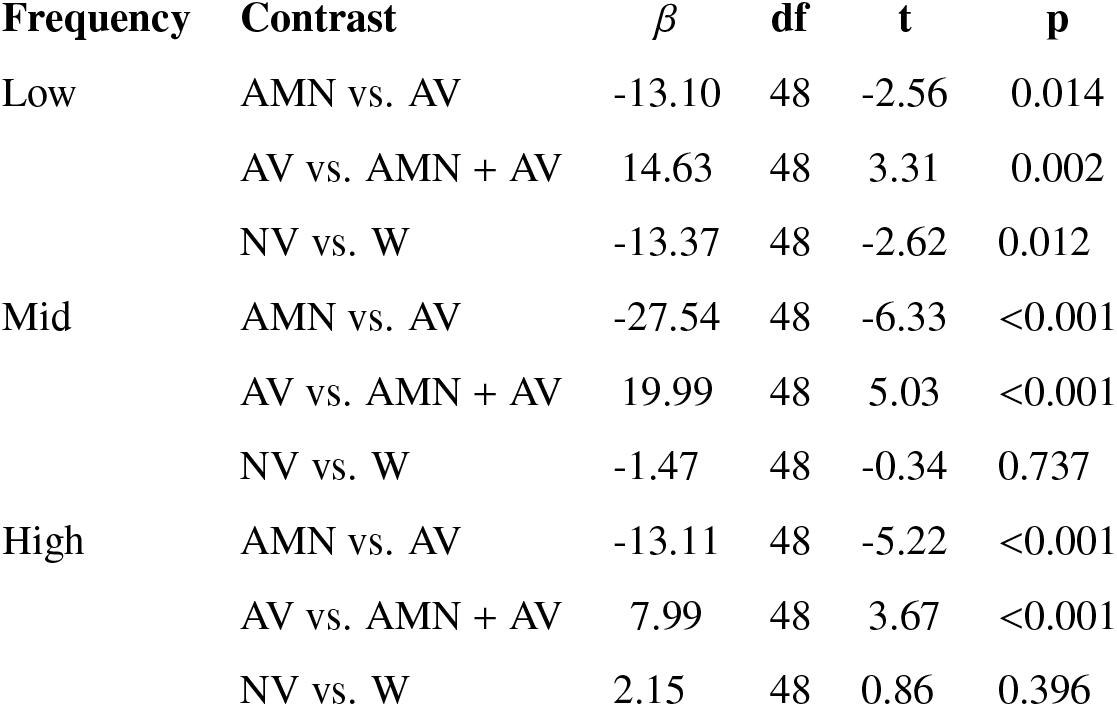
The effect of stimulus complexity per stimulus frequency for response amplitude

| Frequency | Contrast | $\beta$ | df | t | p |
| --- | --- | --- | --- | --- | --- |
| Low | AMN vs. AV | -13.10 | 48 | -2.56 | 0.014 |
|  | AV vs. AMN + AV | 14.63 | 48 | 3.31 | 0.002 |
|  | NV vs. W | -13.37 | 48 | -2.62 | 0.012 |
| Mid | AMN vs. AV | -27.54 | 48 | -6.33 | <0.001 |
|  | AV vs. AMN + AV | 19.99 | 48 | 5.03 | <0.001 |
|  | NV vs. W | -1.47 | 48 | -0.34 | 0.737 |
| High | AMN vs. AV | -13.11 | 48 | -5.22 | <0.001 |
|  | AV vs. AMN + AV | 7.99 | 48 | 3.67 | <0.001 |
|  | NV vs. W | 2.15 | 48 | 0.86 | 0.396 |

**Table A.2:** The effect of stimulus complexity per stimulus frequency for noise amplitude

| Frequency | Contrast | $\beta$ | df | t | p |
| --- | --- | --- | --- | --- | --- |
| Low | AMN vs. AV | 0.48 | 48 | 0.47 | 0.641 |
|  | AV vs. AMN + AV | 0.41 | 48 | 0.47 | 0.644 |
|  | NV vs. W | -1.21 | 48 | -1.19 | 0.241 |
| Mid | AMN vs. AV | -0.19 | 48 | -0.49 | 0.629 |
|  | AV vs. AMN + AV | 0.43 | 48 | 1.27 | 0.209 |
|  | NV vs. W | -0.72 | 48 | -1.84 | 0.072 |
| High | AMN vs. AV | -0.23 | 48 | -0.79 | 0.434 |
|  | AV vs. AMN + AV | 0.48 | 48 | 1.89 | 0.065 |
|  | NV vs. W | -0.08 | 48 | -0.28 | 0.781 |

**Table A.3:**
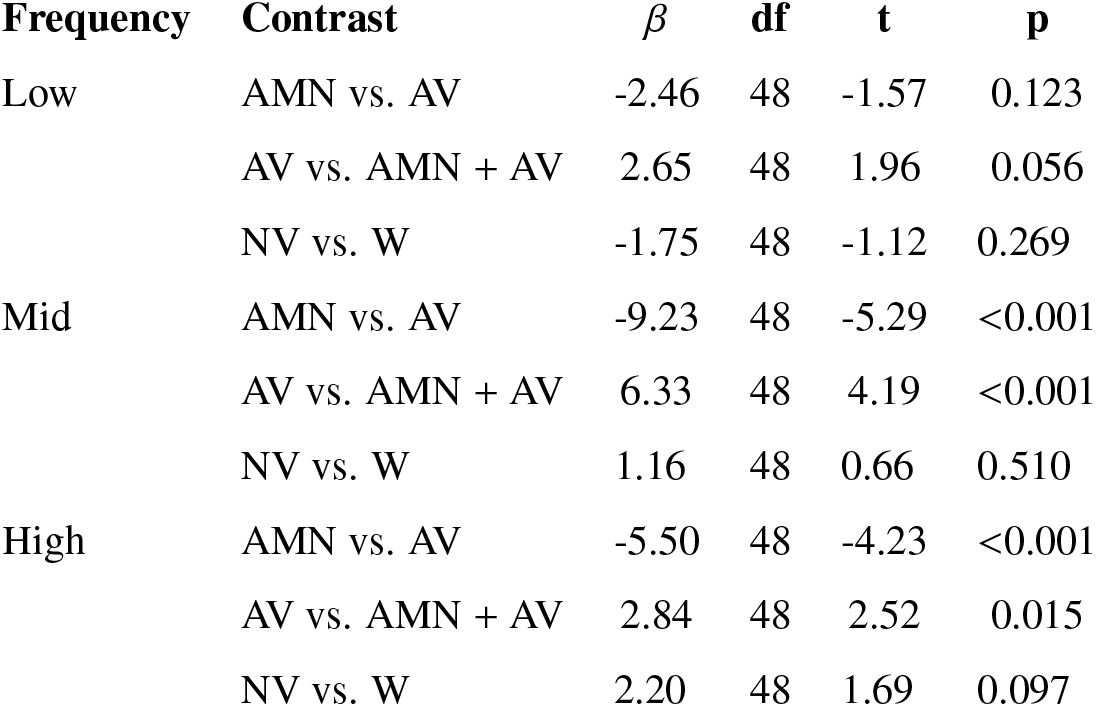
The effect of stimulus complexity per stimulus frequency for response SNR

| Frequency | Contrast | $\beta$ | df | t | p |
| --- | --- | --- | --- | --- | --- |
| Low | AMN vs. AV | -2.46 | 48 | -1.57 | 0.123 |
|  | AV vs. AMN + AV | 2.65 | 48 | 1.96 | 0.056 |
|  | NV vs. W | -1.75 | 48 | -1.12 | 0.269 |
| Mid | AMN vs. AV | -9.23 | 48 | -5.29 | <0.001 |
|  | AV vs. AMN + AV | 6.33 | 48 | 4.19 | <0.001 |
|  | NV vs. W | 1.16 | 48 | 0.66 | 0.510 |
| High | AMN vs. AV | -5.50 | 48 | -4.23 | <0.001 |
|  | AV vs. AMN + AV | 2.84 | 48 | 2.52 | 0.015 |
|  | NV vs. W | 2.20 | 48 | 1.69 | 0.097 |

### A.2. The effect of stimulus frequency per stimulus complexity

**Figure A.2:**
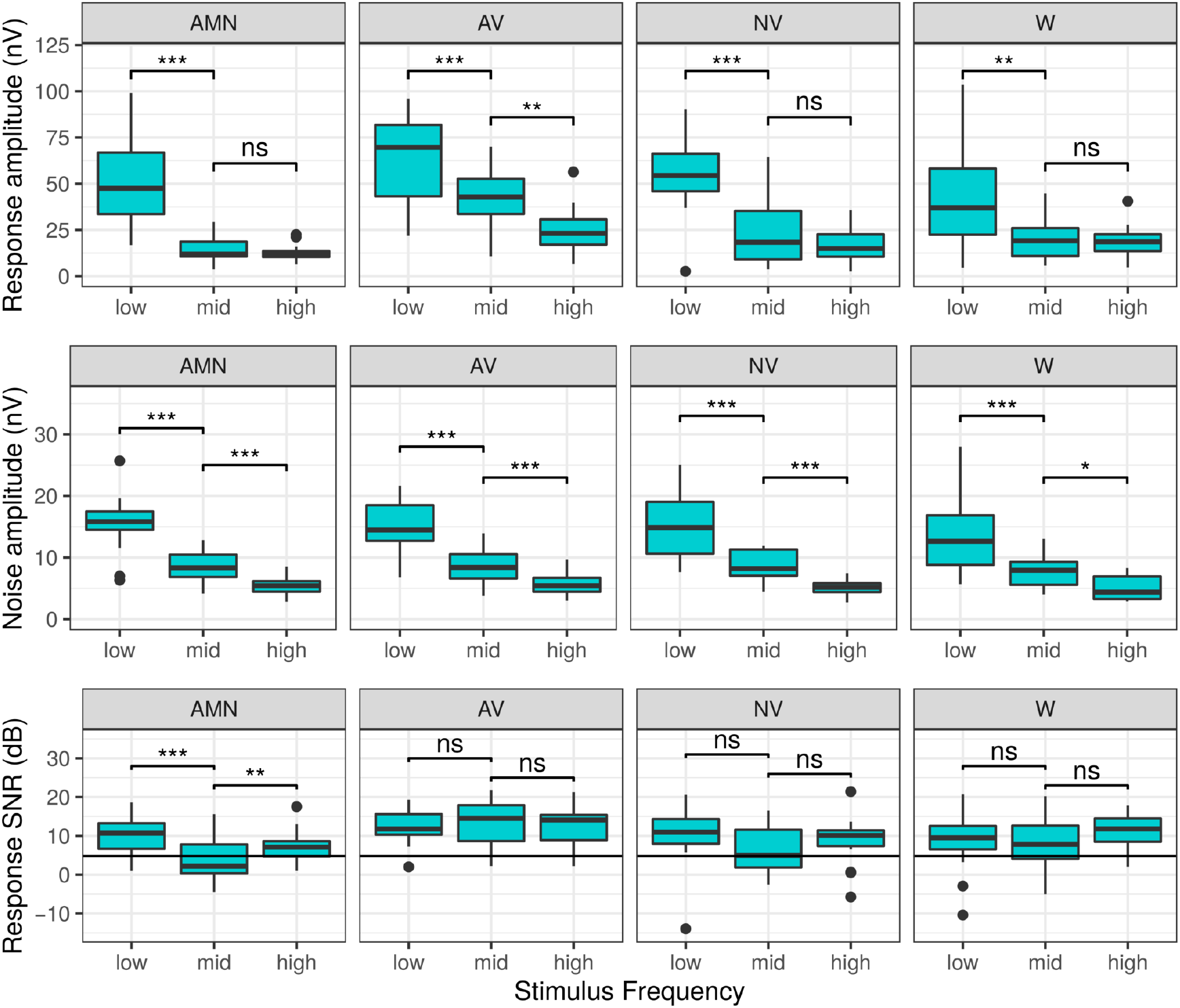
The effect of stimulus frequency on response amplitude, noise amplitude and SNR per stimulus complexity. AMN = amplitude modulated noise, AV = artificial vowel, NV = natural vowel, W = word. Significance codes: *** < 0.001, ** < 0.01, * < 0.05, ns = non significant.

**Table A.4:** The effect of stimulus frequency per stimulus complexity for response amplitude

| **Frequency** | **Contrast** | $\beta$ | **df** | **t** | **p** |
| --- | --- | --- | --- | --- | --- |
| AMN | low vs. mid | -35.62 | 32 | -7.99 | <0.001 |
|  | mid vs. high | 1.99 | 32 | 0.45 | 0.659 |
| AV | low vs. mid | -21.19 | 32 | -3.91 | <0.001 |
|  | mid vs. high | 16.42 | 32 | 3.03 | 0.005 |
| NV | low vs. mid | -32.50 | 32 | -6.19 | <0.001 |
|  | mid vs. high | 6.23 | 32 | 1.19 | 0.245 |
| W | low vs. mid | -20.60 | 32 | -3.46 | 0.001 |
|  | mid vs. high | 2.61 | 32 | 0.44 | 0.664 |

**Table A.5:** The effect of stimulus frequency per stimulus complexity for noise amplitude

| **Frequency** | **Contrast** | $\beta$ | **df** | **t** | **p** |
| --- | --- | --- | --- | --- | --- |
| AMN | low vs. mid | -6.95 | 32 | -9.18 | <0.001 |
|  | mid vs. high | 3.14 | 32 | 4.14 | <0.001 |
| AV | low vs. mid | -6.29 | 32 | -7.98 | <0.001 |
|  | mid vs. high | 3.10 | 32 | 3.93 | <0.001 |
| NV | low vs. mid | -6.55 | 32 | -7.60 | <0.001 |
|  | mid vs. high | 3.46 | 32 | 4.01 | <0.001 |
| W | low vs. mid | -6.06 | 32 | -5.83 | <0.001 |
|  | mid vs. high | 2.83 | 32 | 2.72 | 0.01 |

**Table A.6:** The effect of stimulus frequency per stimulus complexity for response SNR

| **Frequency** | **Contrast** | $\beta$ | **df** | **t** | **p** |
| --- | --- | --- | --- | --- | --- |
| AMN | low vs. mid | -5.62 | 32 | -5.36 | <0.001 |
|  | mid vs. high | -3.18 | 32 | -3.03 | 0.005 |
| AV | low vs. mid | 1.15 | 32 | 0.86 | 0.398 |
|  | mid vs. high | 0.55 | 32 | 0.41 | 0.683 |
| NV | low vs. mid | -3.98 | 32 | -1.70 | 0.096 |
|  | mid vs. high | -2.41 | 32 | -1.03 | 0.308 |
| W | low vs. mid | -1.07 | 32 | -0.50 | 0.621 |
|  | mid vs. high | 2.39 | 32 | 1.12 | 0.272 |

### A.3. The effect of frequency contour per stimulus complexity

**Figure A.3:**
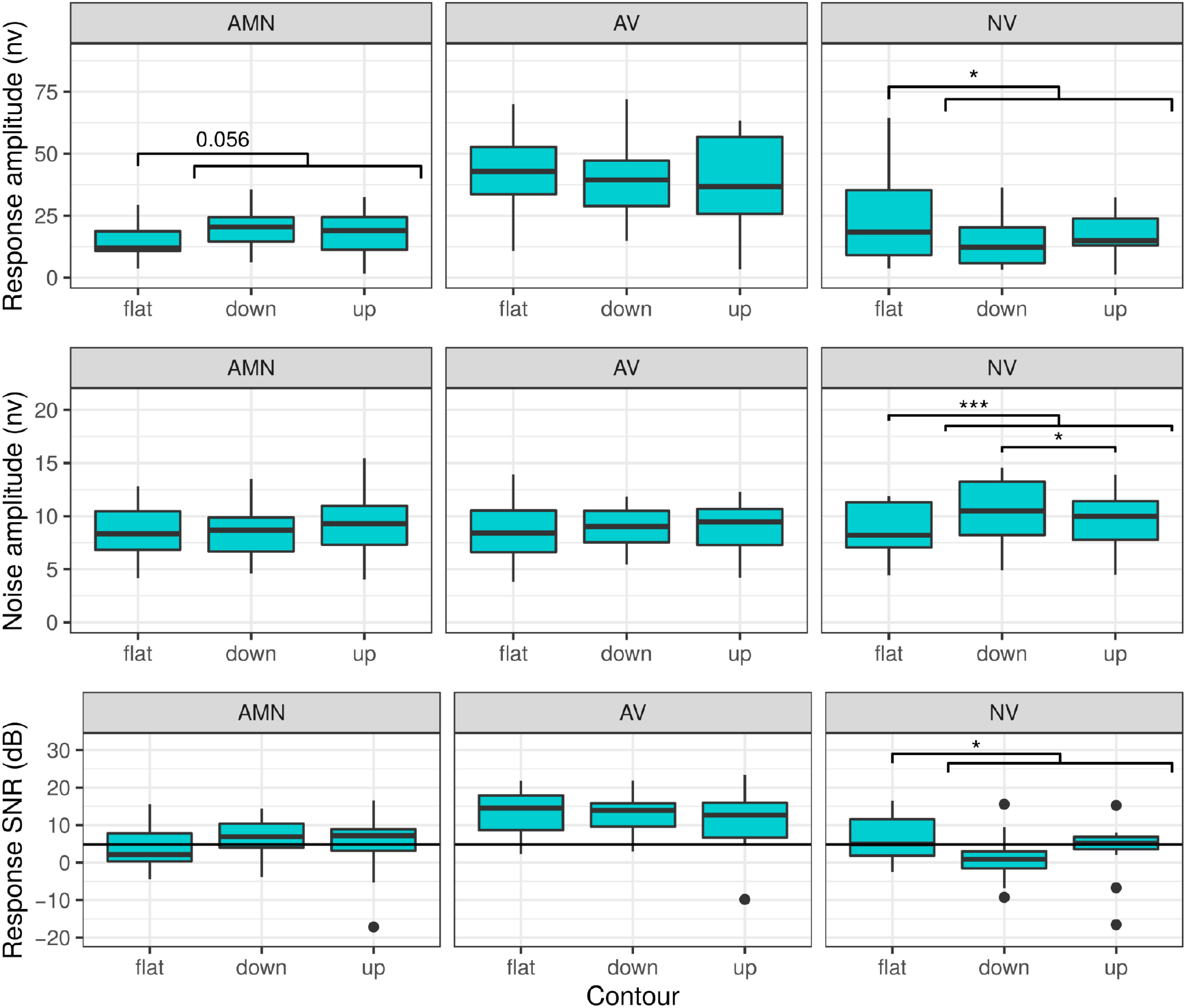
The effect of stimulus contour on response amplitude, noise amplitude and SNR per stimulus complexity. AMN = amplitude modulated noise, AV = artificial vowel, NV = natural vowel, W = word. Significance codes: *** < 0.001, ** < 0.01, * < 0.05, ns = non significant.

**Table A.7:**
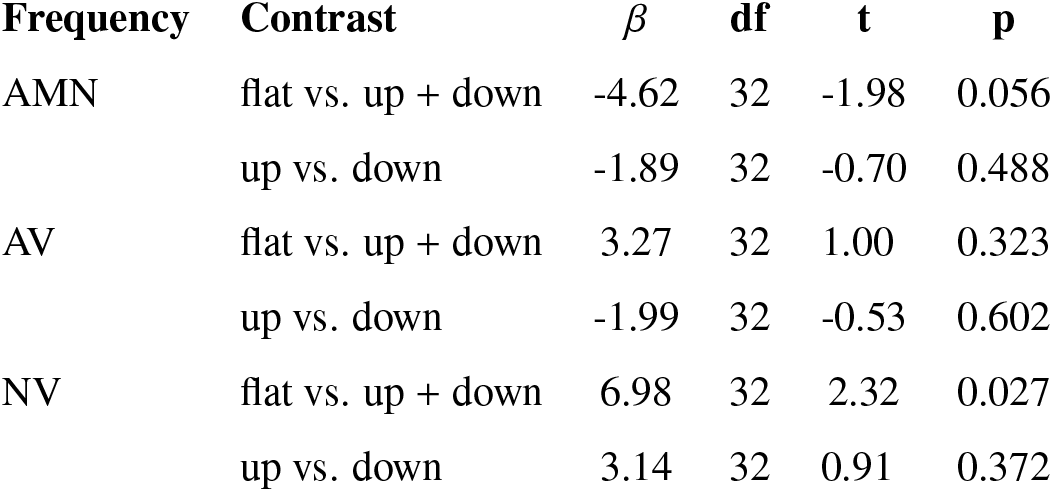
The effect of frequency contour per stimulus complexity for response amplitude

**Table A.8:** The effect of frequency contour per stimulus complexity for noise amplitude

| **Frequency** | **Contrast** | $\beta$ | **df** | **t** | **p** |
| --- | --- | --- | --- | --- | --- |
| AMN | flat vs. up + down | -0.44 | 32 | -1.21 | 0.236 |
|  | up vs. down | 0.79 | 32 | 1.89 | 0.068 |
| AV | flat vs. up + down | -0.12 | 32 | -0.63 | 0.532 |
|  | up vs. down | -0.02 | 32 | -0.04 | 0.967 |
| NV | flat vs. up + down | -1.39 | 32 | -4.54 | <0.001 |
|  | up vs. down | -0.83 | 32 | -2.34 | 0.026 |

**Table A.9:**
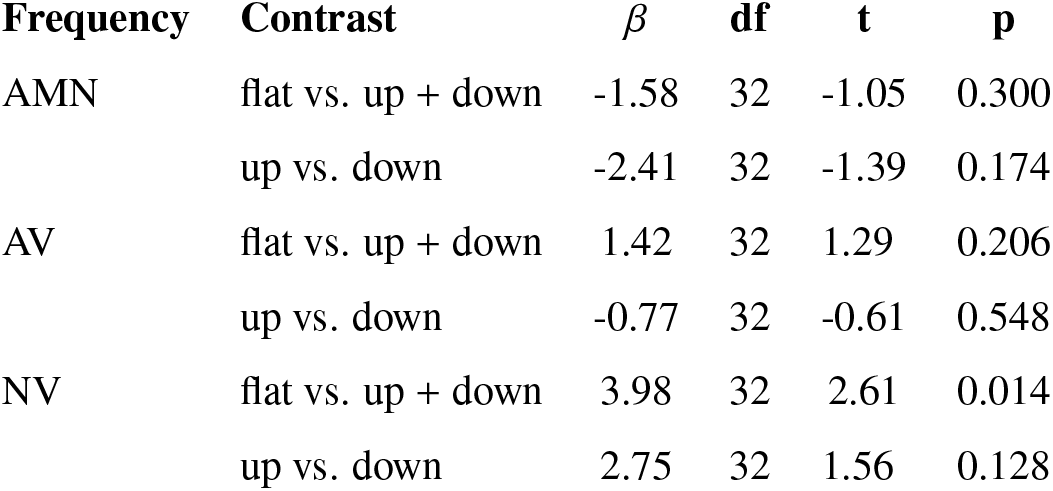
The effect of frequency contour per stimulus complexity for response SNR

| **Frequency** | **Contrast** | $\beta$ | **df** | **t** | **p** |
| --- | --- | --- | --- | --- | --- |
| AMN | flat vs. up + down | -1.58 | 32 | -1.05 | 0.300 |
|  | up vs. down | -2.41 | 32 | -1.39 | 0.174 |
| AV | flat vs. up + down | 1.42 | 32 | 1.29 | 0.206 |
|  | up vs. down | -0.77 | 32 | -0.61 | 0.548 |
| NV | flat vs. up + down | 3.98 | 32 | 2.61 | 0.014 |
|  | up vs. down | 2.75 | 32 | 1.56 | 0.128 |

## Acknowledgements and Funding

Authors would like to thank all subjects who participated in the study. We also want to acknowledge Maud Lobel for her help with data collection and Lotte Cools for lending her voice for stimulus recording. This research was funded by TBM-project LUISTER (T002216N) from the Research Foundation Flanders (FWO) and also jointly by Cochlear Ltd. and Flanders Innovation & Entrepreneurship (formerly IWT), project 50432. Additionally, this project has received funding from the European Research Council under the European Unions Horizon 2020 research and innovation programme (grant agreement No. 637424, ERC starting grant to Tom Francart). The first author, Jana Van Canneyt, is supported by a PhD grant for Strategic Basic research by the Research Foundation Flanders (FWO), project number 1S83618N. There are no conflicts of interest, financial, or otherwise.

## Abbreviations

AM: amplitude modulation
AMN: amplitude modulated noise
AV: artificial vowel
EEG: electroencephalogram
f0: fundamental frequency of the voice
FFR: frequency following response
LMM: linear mixed model
NV: natural vowel
SNR: signal-to-noise ratio
W: word

